# Strong sequencing bias in Nanopore and PacBio prevents assembly of *Drosophila melanogaster* Y-linked genes

**DOI:** 10.1101/2025.02.23.639762

**Authors:** A. Bernardo Carvalho, Bernard Y. Kim, Fabiana Uno

## Abstract

Nanopore and PacBio are generally considered free from sequence composition bias, a key factor – alongside read length – that explains their success in producing high quality genome assemblies. However, our study reveals a systematic failure of both technologies to sequence and assemble specific exons of *Drosophila melanogaster* genes, indicating an overlooked limitation. Namely, multiple Y-linked exons are nearly or completely absent from raw reads produced by deep sequencing with state-of-the-art Nanopore (10.4 flow cells, 200× coverage) and PacBio (HiFi 50×). The same exons are accurately assembled using Illumina 65× coverage. We found that these missing exons are consistently located near simple satellite sequences, where sequencing fails at multiple levels: read initiation (very few reads start within satellite regions), read elongation (satellite-containing reads are shorter on average), and base-calling (quality scores drop as sequencing enters a satellite sequence). These findings challenge the assumption that long-read technologies is unbiased and reveal a critical barrier to assembling sequences near repetitive regions. As large-scale sequencing projects move towards telomere-to-telomere assemblies in a wide range of organisms, recognizing and addressing these biases will be important to achieving truly complete and accurate genomes. Additionally, the underrepresented Y-linked exons provides a valuable benchmark for refining those sequencing technologies while improving the assembly of the highly heterochromatic and often neglected *Drosophila* Y chromosome.

## Introduction

Third-generation sequencing technologies (“TGS”) PacBio and Nanopore are revolutionizing genomics: they are making chromosome level assemblies routine, and full diploid, telomere-to-telomere assemblies (T2T) are becoming the standard for human genome assemblies. TGS employs single-molecule sequencing, avoiding many biases, errors and limitations introduced by cloning or PCR amplification, which affected previous technologies such as Sanger sequencing, Illumina, and 454. A key advantage of TGS is the ability to generate long reads, ranging from approximately 20 kb to over 1 Mb. While initial versions of TGS had high error rates (12% to 20%), the latest error rates are in the range of 0.1% (PacBio HiFi) to 0.5% (Nanopore Q20). Finally, TGS is believed to be nearly bias-free in terms of sequence composition (*e.g.*, Ross et al. 2013), although a few exceptions have been noted. Nurk et al. (2020; 2022) reported a lack of PacBio HiFi sequencing coverage across GA-rich sequences found at a few sites in the human genome, and Flynn et al. (2020) found that no current technology (Illumina, PacBio, or Nanopore) was able to recover the ∼100 Mbp simple satellites (mostly AAACTAT) predicted to occur in the *D. virilis* genome (Gall and Atherton 1974; Bosco et al. 2007). Despite these exceptions, TGS seems to largely fulfill the requirements of Gene Myers’ “perfect assembly” theorem, which he informally stated in tweet format: “Thm: Perfect assembly possible iff a) errors random b) sampling is Poisson c) reads long enough 2 solve repeats.” (Myers 2014). The success of TGS in many organisms attests to this being generally true.

However, when the first TGS dataset of *D. melanogaster* was released (Kim et al. 2014), Carvalho and colleagues (2016) reported an unexpected failure: several single-copy exons of the Y chromosome were either missing or severely underrepresented in the PacBio CLR raw reads, and were missing in the assemblies. This was even more surprising given that the same dataset resolved two challenging Y chromosome regions (Carvalho et al. 2015; Krsticevic et al. 2015). Carvalho et al. (2016) attributed this bias to the use of cesium chloride DNA purification in Kim et al. (2014), which separates DNA based on density. Since the light AT-rich sequences form a band separated from the main DNA band (“satellite bands”; see Altemose 2022, and references therein), and since many *Drosophila* Y-linked introns contain AT-rich satellite DNA (Reugels et al. 2000), they proposed that the missing exons were inadvertently discarded along with these satellite-rich fractions. At the time, the reported sequencing bias seemed to be caused by a fortuitous artifact in sample preparation.

Here we report that the previously observed bias was not caused by cesium chloride purification but is a systematic issue present in all current TGS datasets, regardless of the DNA extraction method. Namely, deep sequencing with state-of-the-art TGS platforms (Nanopore Q20 200× coverage; Kim et al. 2024; PacBio HiFi 50× coverage; Shukla et al. 2024) completely fails to assemble several protein-coding exons of *D. melanogaster* Y-linked genes, whereas Illumina at 65x coverage faithfully assembles these same exons. Further investigation showed that these exons are nearly or completely absent from the raw reads, and are consistently located near simple satellite sequences ( *e.g.*, (AATATAT)_n_ ). We found that these satellites disrupt TGS sequencing at multiple steps: (i) read initiation is inhibited, resulting in very few reads starting within satellite regions; (ii) read elongation is impaired, leading to shorter reads when satellites are present; and (iii) base-calling accuracy declines abruptly when satellite sequences begin, lowering read quality scores. The first two factors (read initiation and extension) result in very low (or zero) coverage of the affected exons in TGS raw reads.

This strong sequencing bias is interesting not only as a phenomenon *per se* but also for its implications in genome assembly and sequencing technology development. While large, highly homogeneous blocks of simple satellites do not occur in humans (see Discussion) and may be rare in most model organisms, the strong bias reported here is relevant to ongoing large-scale sequencing projects such as the Darwin Tree of Life Consortium (2022). The “missing exons” of the *Drosophila* Y-linked genes may also provide a simple, effective and biologically relevant benchmark for improvements in sequencing technology.

## Results

### All TGS technologies fail in the same regions of the *D. melanogaster* Y chromosome

Kim et al. (2024) recently published an extensive Nanopore sequencing dataset of many *Drosophila* species, including a 400× coverage of the reference *D. melanogaster* strain (iso-1). The *D. melanogaster* dataset was generated using Nanopore Q20 (10.4 flow cells), which yields a ∼1% error rate. As DNA was not purified with cesium chloride centrifugation, we expected that the previously observed sequencing bias against Y-linked exons would be absent, *i.e.*, that the *D. melanogaster* Y-linked genes would have uniform coverage (at ∼ 200× ; Kim et al. (2024) used males). However, as shown in Fig. 1 (rows 1 and 2), this was not the case; indeed, the coverage profile in the Nanopore Q20 dataset was essentially identical to that observed in the earlier PacBio CLR dataset (Kim et al. 2014).

**Figure 1.**
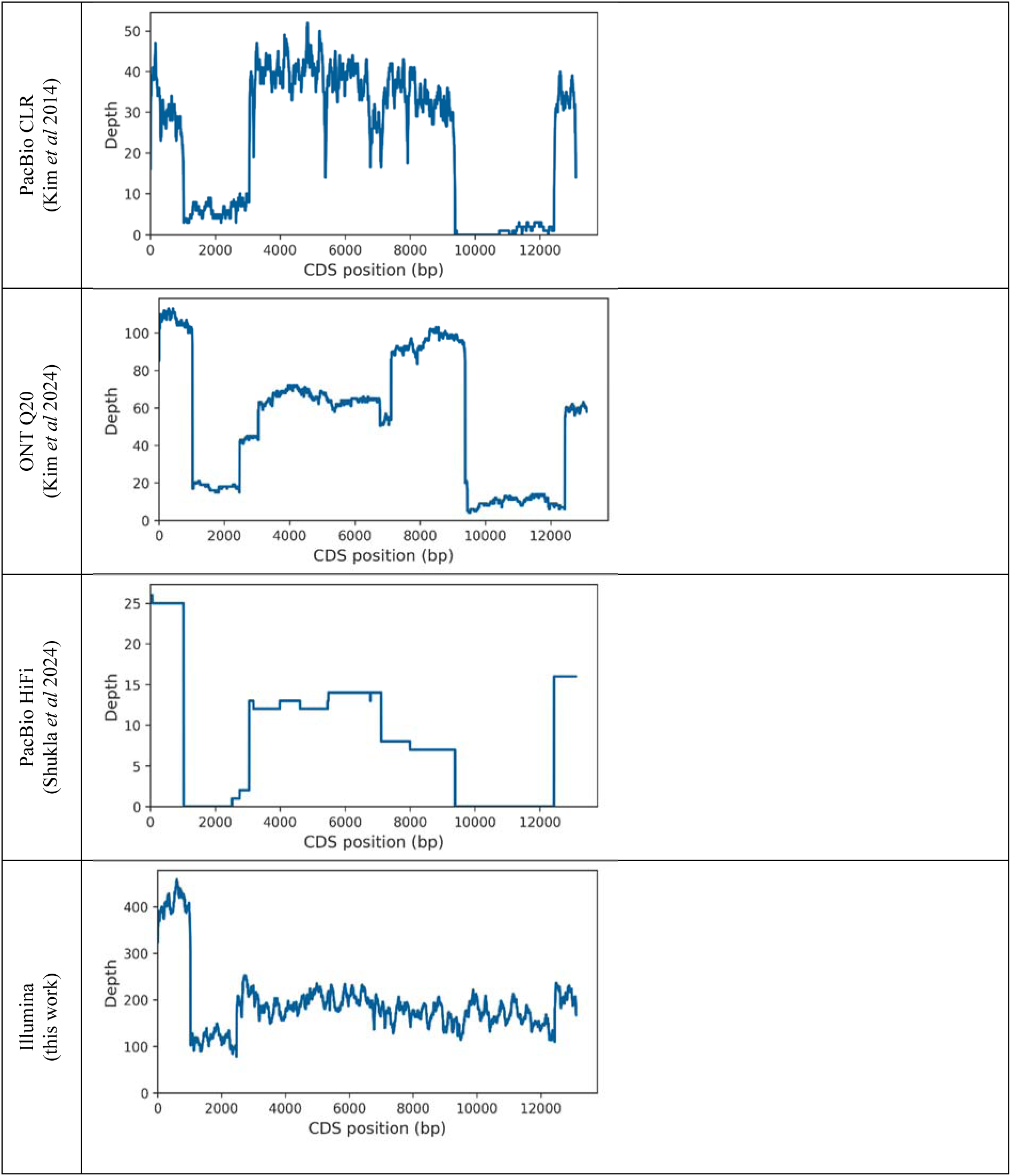
Coverage of the Y-linked gene *kl-3* by raw reads across different sequencing technologies. The three TGS datasets exhibit similar coverage profiles, with two regions of the *kl-3* coding sequence (CDS) nearly absent in the raw reads. In contrast, these regions are well represented in the Illumina reads. See Supplemental Fig. S2 for a similar analysis of the other Y-linked genes.

Furthermore, males of the same strain were sequenced at 100× coverage (50× for the Y chromosome) using PacBio HiFi by (Shukla et al. 2024), again without cesium chloride purification. Nevertheless, the coverage profile was essentially the same, further reinforcing the conclusion that the observed sequencing bias is not linked to the DNA purification method. As a control, we sequenced iso1 males with Illumina at high coverage (∼550×) and found that all previously missing exons were faithfully represented in the raw reads, confirming the earlier results obtained by Carvalho et al. (2016) using a smaller Illumina dataset.

As before, the only major “anomalies” in Illumina coverage are occasional exon duplications (*e.g.*, the initial exons of *kl-3*, visible in Fig. 1), which are fairly common in heterochromatic regions (*e.g.*, Tobler et al. 2017; Chang and Larracuente 2019). As expected, the assembly of these strongly biased TGS datasets consistently failed to assemble many exons, whereas the Illumina assembly recovered nearly all exons (Supplemental Fig. S1). All datasets yielded fragmented assemblies as some Y-linked introns are extremely long (in the Mbp range) and filled with repetitive DNA (Carvalho et al. 2000; Reugels et al. 2000), which leads to assembly breaks. As expected, the Illumina assembly is much more fragmented due to the short read length.

Fig. 1 highlights the coverage inconsistency for the *kl-3* gene; coverage of Y-linked exons is irregular in most genes (Supplemental Fig. S2), and several exons are strongly underrepresented or entirely absent (“missing exons”) in multiple Y-linked genes (Fig. 2). On the other hand, a few Y-linked genes (*e.g.*, *Pp1-Y1* and *Pp1-Y2*) display uniform and high coverage in the raw reads, showing that not all Y-linked exons are equally affected. To further assess the coverage uniformity, we looked at a random sample of five autosomal genes and 20 X-linked genes, and found that all have a uniform coverage profile at the expected depth (400× for the autosomal genes, 200× for X-linked ones; data not shown). To summarize, most or all X-linked and autosomal genes and some Y-linked genes are represented in TGS reads as expected by a random (Poisson) sampling of the genome, whereas most Y-linked genes show highly irregular coverage across different exons, with some exons being nearly absent. Notably, this bias is not exclusive to the Nanopore dataset, as these observations hold for the other TGS datasets mentioned above (PacBio CLR and PacBio HiFi).

**Figure 2.**
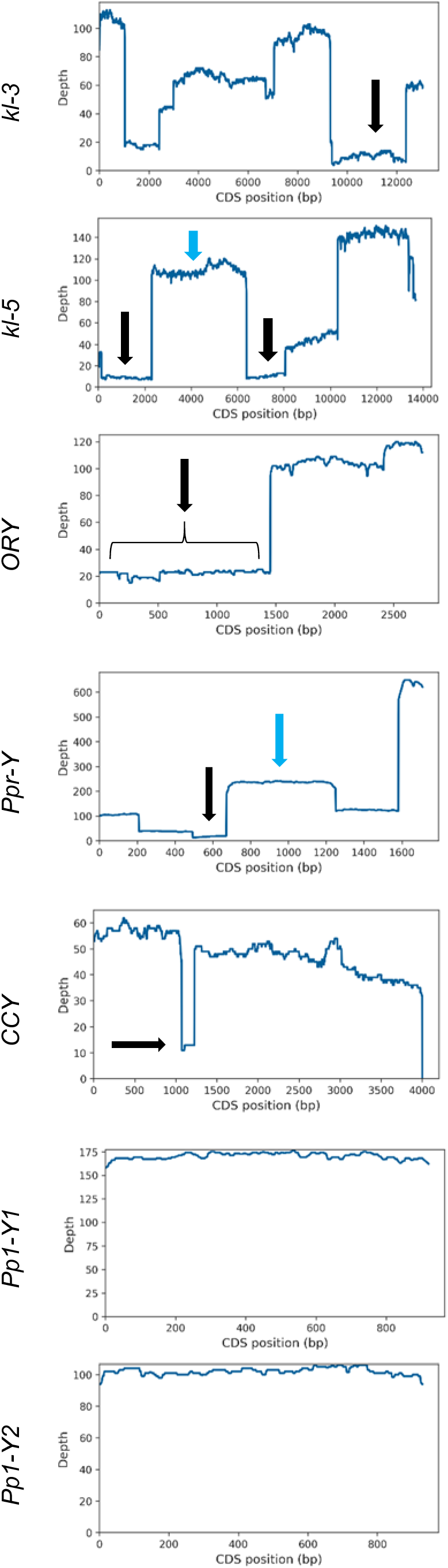
Coverage of selected Y-linked genes in Nanopore raw reads (Kim et al. 2024). *Pp1-Y1* and *Pp1-Y2* exhibit uniform coverage, contrasting with other Y-linked genes with highly irregular coverage. Arrows point to the missing exons (black arrows) and normal coverage exons (blue arrows) investigated in detail. See Supplemental Fig. S2 for the coverage data on additional Y-linked genes.

Finally, even the *Pp1-Y1* and *Pp1-Y2* genes, which have uniform and high coverage (160× and 100×, respectively), are below the expected coverage of ∼200×. This discrepancy is probably caused by polytenic tissues in adult males, where heterochromatic regions are known to be severely undereplicated (Yarosh and Spradling 2014). In line with this, Flynn et al. (2020) observed that *D. virilis* adult whole flies contain 40% fewer heterochromatic satellites than fully diploid tissues, strongly suggesting that the underrepresentation of heterochromatin in polytenic tissues systematically lowers sequencing coverage of these regions in adult flies. This phenomenon differs from the “missing exons” we are investigating here, and does not cause assembly problems.

### A detailed investigation of the missing exons

Our results show that certain regions of the *D. melanogaster* Y chromosome are recalcitrant to TGS. But what exactly is inside these regions? To investigate this, we used the Nanopore Q20 dataset from Kim et al. (2024), as its huge coverage allowed us to recover a small number of surviving reads from the missing exons. Given the consistency of the bias across platforms, we presume that the same phenomenon is happening with the shallower PacBio CLR and PacBio HiFi datasets (Kim et al. 2014; Shukla et al. 2024). First, we used a simple blastN search to recover the few reads covering the missing exons (black arrows in Fig. 2) and assembled them using a variety of approaches. Commonly used tools such as Canu (Koren et al. 2017) and Flye (Kolmogorov et al. 2019) either failed entirely or yielded very small contigs. The most effective approach we found was to tweak several parameters of *miniasm* (Li 2016); this yields draft assemblies of all missing exons, with contigs sizes ranging from 12 kb to 102 kb (Methods). With these draft assemblies, we could now look at what is inside these regions. As shown in Fig. 3, all Y-linked exons, both those with normal coverage and those with low coverage in TGS datasets, are embedded in highly repetitive DNA (∼ 95% of repetitive DNA). However, there is an important difference between them: exons that are missing or underrepresented in TGS datasets are closely associated with simple satellite sequences ( *e.g.*, (AAGAA)_n_, (AATATA)_n_), whereas exons with normal coverage are devoid (or nearly so) of satellite DNA in their vicinity. Instead, these regions are embedded in transposable elements (TEs). These consistent association patterns suggest that simple satellite DNA is the major factor disrupting TGS sequencing. For the sake of simplicity, we classified the *kl-5* exons 10-12 region as a “normal coverage” region, but it actually is an intermediate case: it has satellite blocks but they are not in close proximity to the exons (Fig. 3), the raw read statistics (discussed below) are only mildly disturbed, and it has fairly high coverage (Fig. 2).

**Figure 3.**
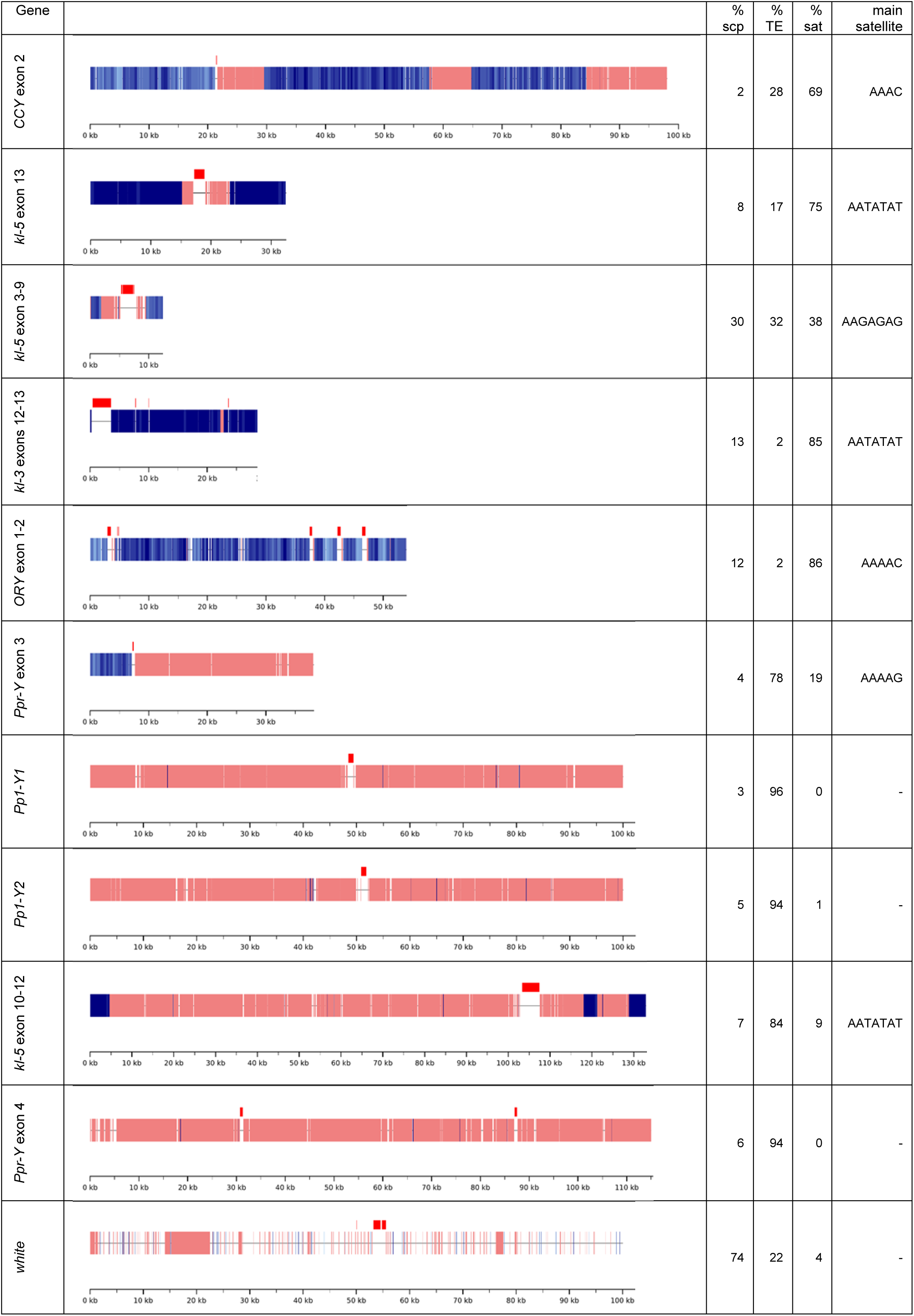
Sequence composition of selected regions of the *D. melanogaster* Y chromosome, and of a control euchromatic region (*white* gene). Red, gene CDS (including pseudogenes); pink, transposable elements; blue, simple satellites (shades of blue are proportional to AT-richness); thin line, single-copy sequences. The top six regions all have very low read coverages, whereas the bottom five have approximatelly normal coverages.

### How does simple satellite DNA interfere with TGS?

A careful examination of the few surviving reads from the missing exons suggested several potential mechanisms underlying this sequencing bias. First, the base quality values drop precipitously as sequencing enters the satellite blocks (Supplemental Fig. S3). Since Nanopore sequencing software filters low-quality reads and put them into a separate file by default, we initially reasoned that the missing Y-linked reads might be found in this low-quality subset. However, after examining these files, we found no enrichment of missing exon reads (data not shown), ruling out read quality as the primary explanation for the missing exons.

We also noticed partial exon duplications (*i.e.*, pseudogenes) occurring in tandem with their functional copies. In some cases, these pseudogenes are in reverse orientation in relation to the their functional copies (*e.g.*, read SRR26246282.1135981 has two copies of *kl-5* exon 13 in reverse orientation). Since single-strand DNA forms at some point in all TGS technologies, these reverse-oriented copies could theoretically generate hairpin structures that might interfere with sequencing. However, this could not be a general explanation since most missing exons lack reverse-oriented pseudogenes. Furthermore, in most cases, the reverse-oriented sequences seem to be sequencing artifacts not present in the genome (Supplemental Fig. S10).

Two additional observations proved to be more fruitful. First, we found that reads from the missing exons were noticeably shorter. Indeed, there is a statistically significant difference in size between them and reads from normal coverage Y-linked regions (Fig. 4). This observation strongly suggests that simple satellite sequences disturb the read traversal across the pores (“read elongation”, for short), either by slowing it or causing a premature termination. Regardless of the precise mechanism, this effect alone would reduce the sequencing coverage in the affected regions, making exons near satellites disproportionally underrepresented. We will deal with the second observation in the next section.

**Figure 4.**
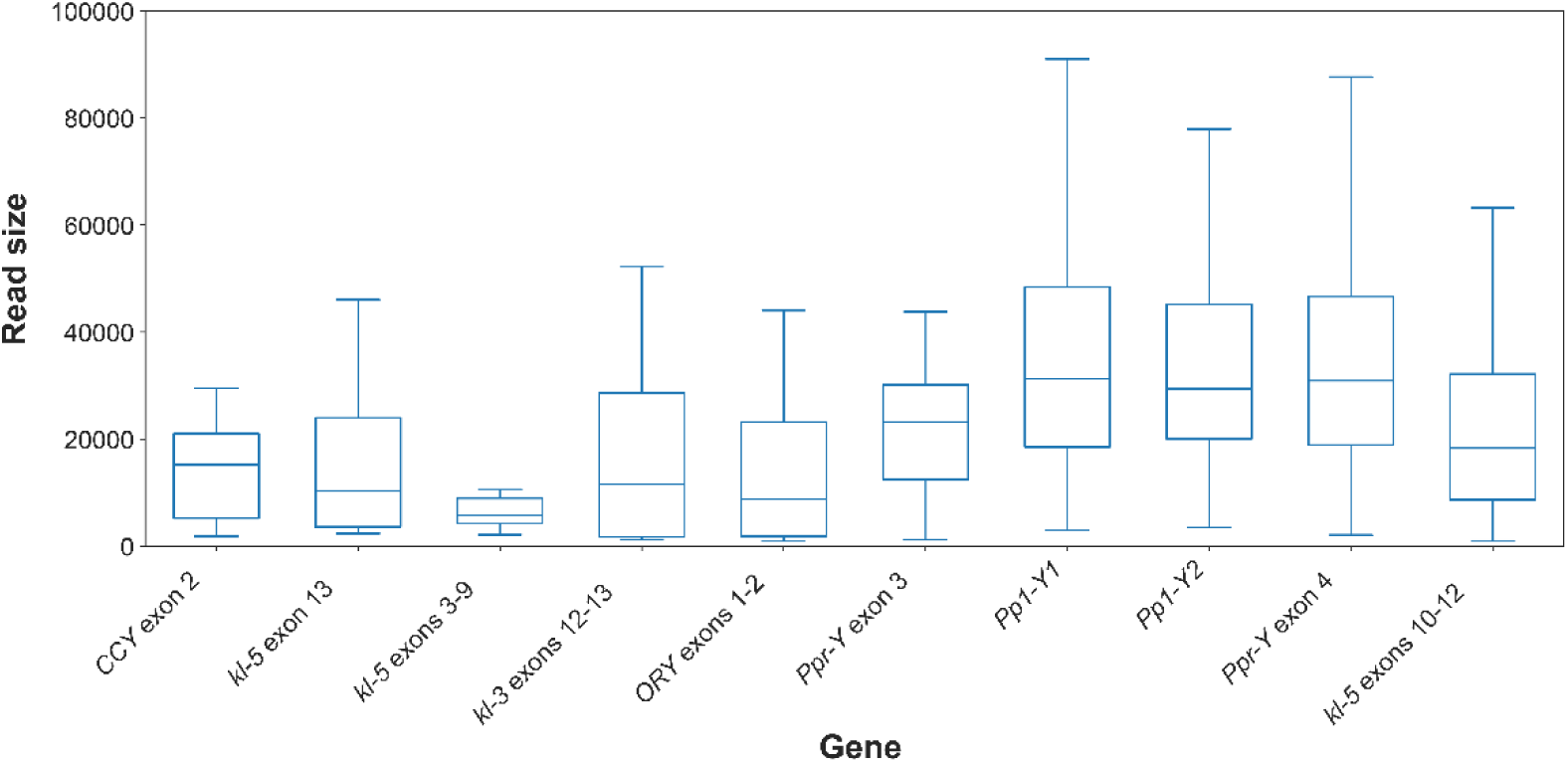
Raw read size in selected regions of the *D. melanogaster* Y chromosome.The six regions in the left have very low read coverages, whereas the last four have approximatelly normal coverages. The difference in read size between the two groups is statistically significant (*F*_1,8_ = 93.2; *P* = 10^-4^), as are the within-group differences (*F*_8,708_ = 8.6; *P* < 10^-5^ ; nested ANOVA on log-transformed values). Note that the *kl-5* exons 10-12 region represents an intermediate case between low-coverage and normal coverage regions, as commented in the previous section. Note also that Fig. 4 probably underestimates the effect of the satellites on read size, since some regions (*e.g.*, *CCY* exon 2 and *Ppr-Y* exon 3) have a large amount of non-satellite sequence (Fig. 3).

### Satellite DNA as a barrier to read initiation

Our second observation is that the reads covering the missing exons do not seem to be randomly distributed, as indicated by two key findings. First, the exon orientation within the reads (forward/reverse) is strongly skewed in most missing exons, deviating from the 50:50 ratio expected by theory and observed in the four Y-linked regions with normal coverage (Table 1, columns 3-4). Given the small sample sizes for the missing exons, which reduces the statistical power of individual tests, we combined the *P* values from all six missing exons using Fisher’s method (Sokal and Rohlf 1995, p. 794). This combined analysis rejects the null hypothesis of random exon orientation (*χ^2^*= 25.10, 12 *d.f.*, *P* = 0.014). In contrast, applying the same procedure to the four normal coverage regions yields a non-significant *P*-value (*χ^2^* = 3.37, 8 *d.f.*, *P* = 0.91), confirming that forward/reverse exon orientation follows random expectations in these regions.

**Table 1.**
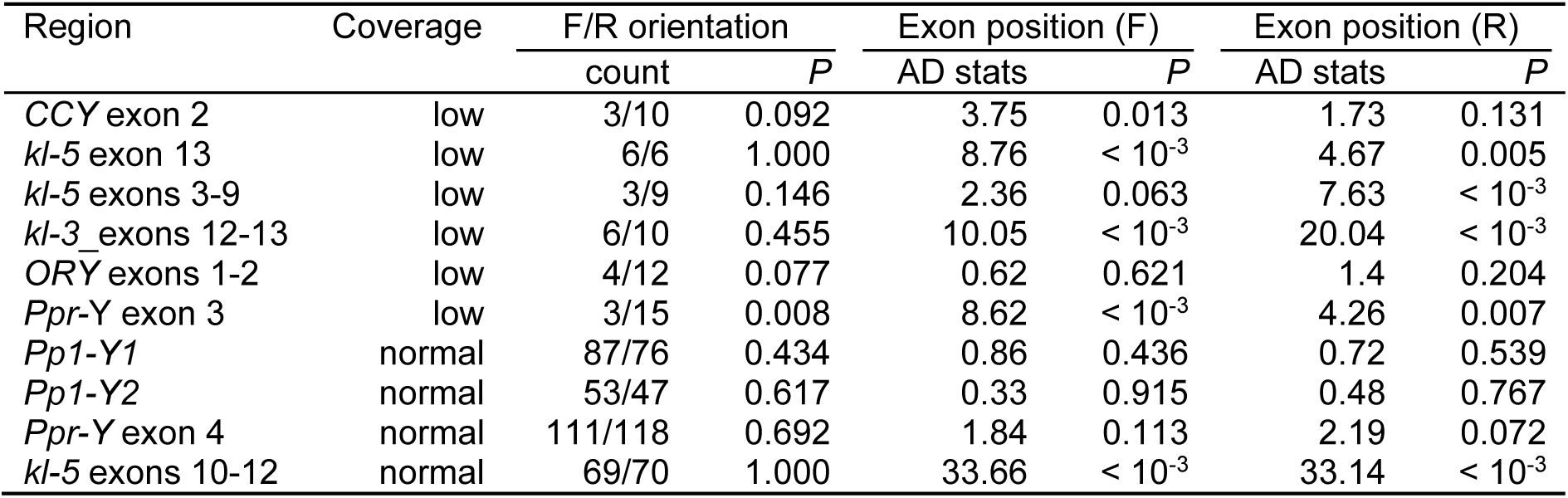
Statistical analysis of exon orientation and position in raw reads. Note that low-coverage regions frequently display skewed exon orientation (F/R; columns 3-4) and non-random exon positions (columns 5-8).

| Region | Coverage | F/R orientation |  | Exon position (F) |  | Exon position (R) |  |
| --- | --- | --- | --- | --- | --- | --- | --- |
|  |  | count | *P* | AD stats | *P* | AD stats | *P* |
| *CCY* exon 2 | low | 3/10 | 0.092 | 3.75 | 0.013 | 1.73 | 0.131 |
| *kl-5* exon 13 | low | 6/6 | 1.000 | 8.76 | $< 10^{-3}$ | 4.67 | 0.005 |
| *kl-5* exons 3-9 | low | 3/9 | 0.146 | 2.36 | 0.063 | 7.63 | $< 10^{-3}$ |
| *kl-3* exons 12-13 | low | 6/10 | 0.455 | 10.05 | $< 10^{-3}$ | 20.04 | $< 10^{-3}$ |
| *ORY* exons 1-2 | low | 4/12 | 0.077 | 0.62 | 0.621 | 1.4 | 0.204 |
| *Ppr-Y* exon 3 | low | 3/15 | 0.008 | 8.62 | $< 10^{-3}$ | 4.26 | 0.007 |
| *Pp1-Y1* | normal | 87/76 | 0.434 | 0.86 | 0.436 | 0.72 | 0.539 |
| *Pp1-Y2* | normal | 53/47 | 0.617 | 0.33 | 0.915 | 0.48 | 0.767 |
| *Ppr-Y* exon 4 | normal | 111/118 | 0.692 | 1.84 | 0.113 | 2.19 | 0.072 |
| *kl-5* exons 10-12 | normal | 69/70 | 1.000 | 33.66 | $< 10^{-3}$ | 33.14 | $< 10^{-3}$ |

A second sign of non-randomness in the missing exons reads is that many of them seem to start at nearly identical genomic positions, instead of being randomly distributed across the entire region of the exon. Supplemental Fig. S4 and Supplemental Fig. S5 illustrate this pattern for the *kl-3* exons 12-13 and the *kl-5* exon 13, respectively, with similar trends observed in most missing exons. We tested this statistically as follows. First, we used blastN to obtain the distance between the start of each read and the target exon (which is the only safe landmark in these repeat-rich regions). If all reads have the same size and read starts are random, the null hypothesis for exon positions would be a simple uniform distribution within the range [1,read size]. However, read sizes are variable and in this case it seems reasonable to suppose that the proper null hypothesis for *n* reads would be a composite of *n* uniform distributions, one for each read size. We confirmed this supposition through simulations, which also validated the statistical test described below (Supplemental Fig. S6 and Supplemental Table S2). We then compared the observed distribution of exon positions against the null hypothesis of random positions, analyzing forward (F) and reverse (R) reads separately due to the previously noted orientation bias. The result using the Anderson-Darling test, which is more sensitive for small sample sizes than the Kolmogorov-Smirnov test (Razali and Yap 2011), is shown in Fig. 5 and Table 1, columns 5-8 (Supplemental Table S3 shows the results for the Kolmogorov-Smirnov test). We found that most “missing exon” regions strongly depart from randomness, whereas reads from the normal coverage regions *Pp1-Y1*, *Pp1-Y2*, and *Ppr-Y* exon 4 followed the expected random distribution of exon positions. The *kl-5* exon 10-12 region is again an exception, but note that its deviations from randomness are mild (Fig. 5; Supplemental Fig. S9). As we commented before, this region has intermediate characteristics between normal and low coverage regions, most likely because it contains satellite DNA, but not in close vicinity to the exons (Fig. 3).

**Figure 5.**
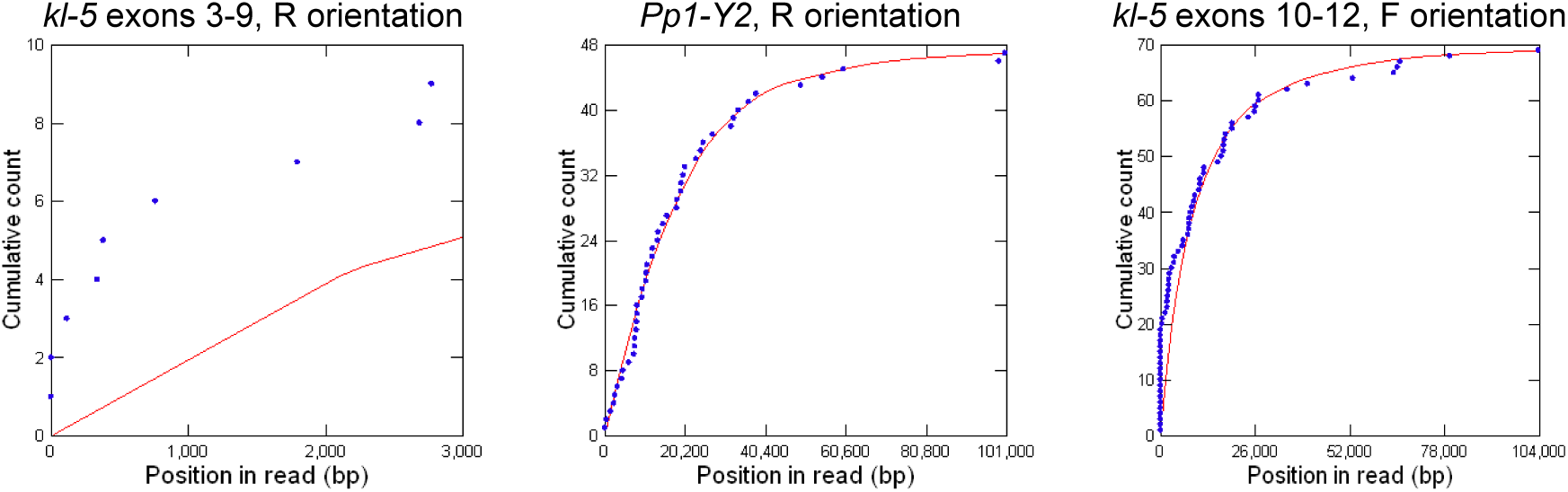
Comparison between observed exon positions within reads (blue dots), and their expected random distributions (red lines). Exon positions are a proxy for read start points in the genome. Left: *kl-5* exons 3-9 region, a low-coverage region (Anderson-Darling test: *P* < 10^-3^); note the huge discrepancy between observed and expected exon positions. Center: *Pp1-Y2*, a normal coverage region (*P* = 0.767); note the very good agreement. Right: *kl-5* exons 10-12, a normal coverage region with intermediate characteristics. Note the fairly good agreement; the *P* value of the Anderson-Darling test (*P* < 10^-3^) reflects the much higher statistical power in normal coverage regions due to their much larger read numbers. See Supplemental Fig. S9 for the remaining regions.

The most likely explanation for this non-randomness both in F/R exon orientation and exon position within reads is that some genomic regions surrounding missing exons have a much lower probability of serving as successful read initiation sites in Nanopore sequencing. As a consequence, read starts would be clustered in some genomic regions and depleted in others, instead of being evenly distributed. Given our previous observations, the most likely culprit for this read initiation suppression is satellite DNA. We tested this hypothesis by counting for each missing exon region how many reads started within satellite blocks and how many started in other sequence types (single-copy, TEs, or the exons themselves). We then compared with a binomial test the observed number of satellite-initiating reads to their expected frequency; the latter is simply the amount of satellite DNA in the region (Fig. 3). As shown in Table 2, there is nearly complete avoidance of satellite DNA as a sequencing starting point. Among the 87 reads covering the missing exons, only one initiated within a satellite block, despite the fact that satellite DNA accounts for 37% to 86% of these regions. This is the main cause of the missing exons: when an exon is located between two large satellite blocks, the only chance of getting sequenced is when reads starts in the small “permissive” regions nearby (TE or single copy), pointing towards it. Reads starting outside the satellite blocks cannot reach the exon because it is too distant, and the satellite DNA cripples the read extension. *Ppr-Y* exon 3 (Fig. 6) illustrates this pattern very well: it is flanked by permissive TEs on the right side and a non-permissive satellite block on the left, and reads only initiate in the permissive regions, never within the satellite DNA.

**Figure 6.**
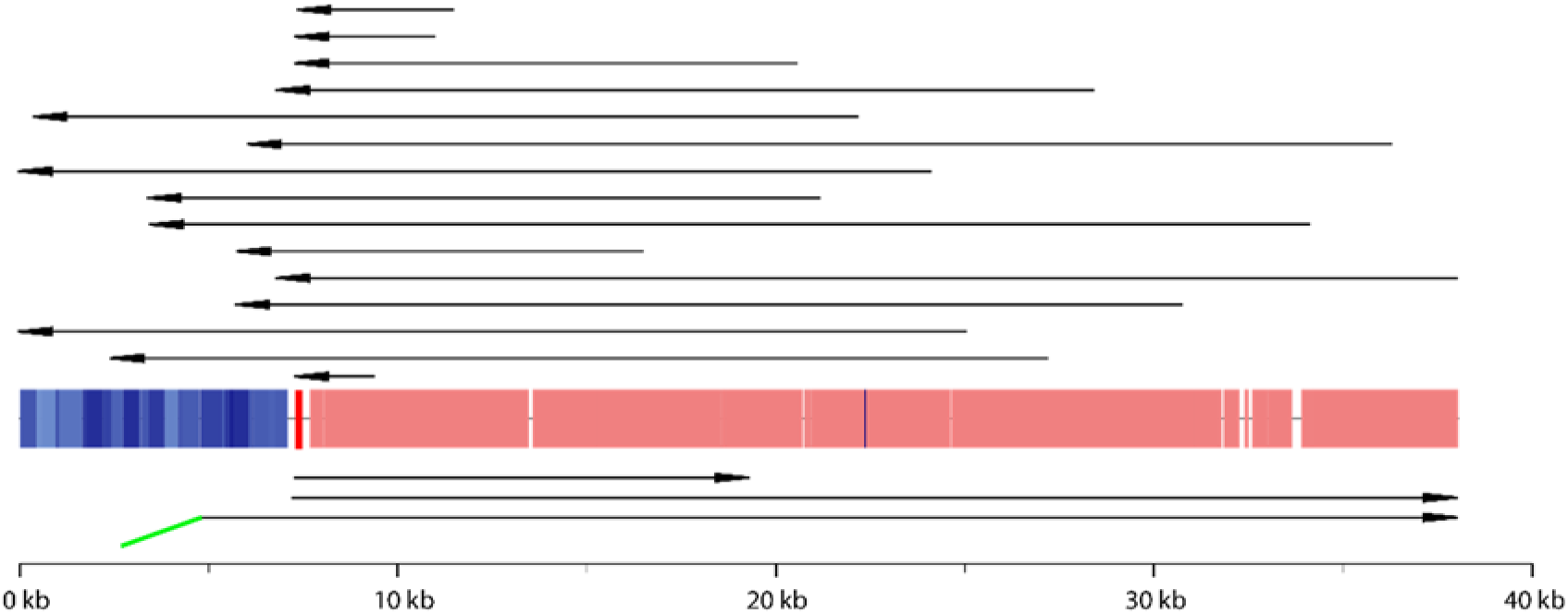
Sequencing bias in the *Ppr-Y* exon 3 region. The region has permissive TEs on the right side (pink) and a non-permissive satellite block on the left (blue). The exon itself (shown in red) is flanked on both sides by tiny single-copy regions (∼190 bp; thin black line). A total of 18 reads cover the exon (black arrows), but their start points are highly asymmetric. Fifteen reads (top) originated in the right side, in scattered points inside the TEs, and extended through the exon and partially into the satellite block. Only two reads originated on the left side, both starting in the tiny 190 bp single-copy region between the satellite block and the exon, extending towards the exon and the TEs. The 18^th^ read (SRR26246282.1589819, shown at the bottom) is an “exception that confirmed the rule”. It starts on the left, seemingly in the middle of the satellite block, but actually is a chimeric read. This read begins with ∼3kb of chromosome 3L (a permissive sequence, shown in green) before transitioning into the end of the satellite block. Note that this avoidance of satellite blocks as sequencing start points explains the low coverage of the exon, the observed exon orientation asymmetry shown in Table 1 (3 forward reads : 15 reverse reads), and the non-random exon position within reads (columns 5-8 of Table 1), since read starts are clustered outside the satellite block.

**Table 2.** Avoidance of satellite DNA as starting points in ONT reads.

| Region | Reads starting in sat. DNA | Reads not starting in sat. DNA | Obs. freq | Exp. freq | *P* |
| --- | --- | --- | --- | --- | --- |
| *CCY* exon 2 | 0 | 13 | 0.0 | 69.2 | $< 10^{-4}$ |
| *kl-5* exons 3-9 | 0 | 12 | 0.0 | 37.5 | 0.005 |
| *kl-5* exon 13 | 0 | 12 | 0.0 | 75.0 | $< 10^{-4}$ |
| *kl-3* exons 12-13 | 0 | 16 | 0.0 | 85.0 | $< 10^{-4}$ |
| *ORY* exons 1-2 | 1 | 15 | 6.2 | 86.5 | $< 10^{-4}$ |
| *Ppr-Y* exon 3 | 0 | 18 | 0.0 | 18.7 | 0.035 |

We found additional chimeric or rearranged reads in other missing exons, suggesting that the scarcity of surviving reads results in an increased proportion of rare sequencing artifacts (Supplemental Fig. S10).

### Are there “missing exons” in non Y-linked *D. melanogaster* genes?

We addressed this question by doing a plain Flye assembly of Kim et al. (2024) Nanopore dataset (Methods) and checking for low CDS coverage (*i.e.*, missing exons) in all *D. melanogaster* genes. This approach detected all Y-linked genes with missing exons discussed here, and two autosomal genes, *Gpa2* and *klhl10* (Larkin et al. 2021), which were completely absent from the Nanopore plain assembly. However, unlike the Y-linked genes, these autosomal genes exhibited uniform raw read coverage with no underrepresented regions (Supplemental Fig. S7). This shows that their assembly failures were due to factors unrelated to sequencing bias. These findings suggest that, in *D. melanogaster*, the close proximity between coding exons and large, homogeneous satellite blocks is restricted to the Y chromosome.

## Discussion

We found that large blocks of simple satellite sequences near several exons of *D. melanogaster* Y-linked genes severely disrupt sequencing with Nanopore and PacBio technologies. This disruption is so strong that the affected exons are barely present in the raw reads and are absent from the final assemblies, even at very high sequencing coverage (*e.g.*, 200×). In contrast, the same exons are faithfully assembled in Illumina datasets, likely because the much shorter fragments employed by this technology (typically 350-600 bp) allow exons to be sampled free from any adjacent satellite DNA. The disruption mentioned above affected even the most complete assembly of the *D. melanogaster* Y available: (Chang and Larracuente 2019), using the shallower and equally biased PacBio CLR dataset from (Kim et al. 2014), had to manually fill the gaps in Y-linked genes by integrating CDS sequences from FlyBase (Larkin et al. 2021).

The bias primarily affects read initiation but also impairs read extension and base calling, with read initiation being the most critical factor. To our knowledge, this is the first systematic and in-depth analysis revealing a severe sequencing bias that affects all TGS platforms. Two previous studies (besides Carvalho et al 2016) have independently detected aspects of this issue in different contexts. Flynn et al. (2020) reported that no current sequencing technology could fully recover the ∼100 Mbp simple satellites estimated to be present in the *D. virilis* genome (Gall and Atherton 1974; Bosco et al. 2007), with PacBio CLR recovering only 10.9 Mbp, Illumina 16.0 Mbp, and Nanopore 28.2 Mbp. They also observed that satellite sequences reduced Illumina read quality scores (other sequencing platforms were not investigated). Similarly, Nurk et al. (2020; 2022) described minor fragmentation in the first T2T human genome assembly due to a lack of PacBio HiFi coverage across GA-rich sequences, suggesting that “this coverage bias appears to be a current weakness of the HiFi chemistry.” While much less detail is available in these two studies, it seems likely that they share the same underlying cause with our study.

Several factors probably contributed to the near absence of prior reports on this bias. First, we could only detect it because the affected Y-linked exons served as sequence landmarks; a missing non-coding sequence in the middle of the Y chromosome would go unnoticed unless someone is attempting a T2T assembly. Second, it affects genes in the Y chromosome, the least known chromosome in *Drosophila*. Third, in the human genome (the only one with extensive T2T assemblies), the bias is much milder and largely restricted to PacBio HiFi, allowing gaps to be closed with Nanopore reads (Nurk et al. 2022).

This mildness initially puzzled us. One possible explanation is the small size of the satellite block – *e.g.*, the chromosome 8 gap identified by Nurk et al. (2022) was caused by a mere 256 bp (AAAGG)_n_ sequence. However, large blocks of simple satellites do occur in the human genome. Namely, HSat2 blocks (monomer: CATTCGATTC) reach up to 12.6 Mbp, and a Hsat3 block (monomer: CATTC) in chromosome 9 has 27.6 Mbp (Altemose et al. 2022). How could these regions be successfully assembled while *Drosophila* Y-linked exons were lost? We believe the key difference is sequence homogeneity. Specifically, these satellite blocks could only be sequenced and assembled because they are much less homogeneous than their *Drosophila* counterparts. As shown in Table 3, the longest perfect tandem repeat within the 27.6 Mpb human hsat3_9_3 block (a CATTC monomer) is only 20 units long. In contrast, despite being much smaller in total length, the unfinished sequences of *Drosophila* shown in Table 3 (actually, raw reads) have hundreds of perfect tandem repeats. The latter number is probably a severe underestimation since any sequencing error would artificially break a perfect repeat block. Another hint that homogeneity (rather than size) is the key problem is that the human chromosome 8 (AAAGG)_n_ sequence mentioned above is a perfect tandem repeat of 51 monomers. Although less detail is available for *D. virilis*, Flynn et al. (2020) estimated its satellite sequence identity at 98.5% to 99% in Illumina reads, lower than what we observe for *D. melanogaster* Y satellites. This heterogeneity might explain why *D. virilis* satellites were partially recovered (28.2 %) in Nanopore reads, a much higher rate than the values observed in our *D. melanogaster* dataset (Fig. 2). Unfortunately, the Nanopore data of *D. virilis* came from the older flow-cells 9.4 with higher error rates ( > 5% ), preventing a more precise analysis.

**Table 3.**
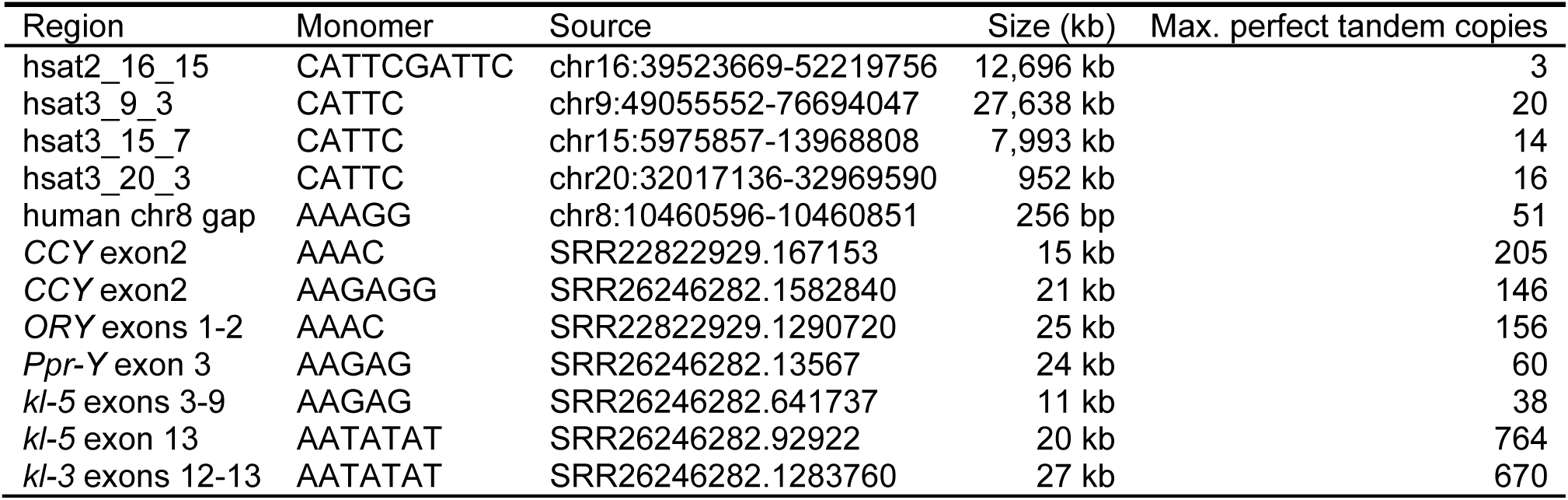
Homogeneity of satellite blocks in the human and *D. melanogaster* genomes.

The above findings strongly suggest that *Drosophila* Y-linked satellite blocks are orders of magnitude more homogeneous than the human satellites (and possibly of *D. virilis* as well), and that the heterogeneity present in the latter prevents or at least attenuates the bias during TGS sequencing. These results also imply that a complete telomere-to-telomere assembly of *D. melanogaster,* including the Y chromosome, will have to await improvements in sample preparation and/or sequencing technology.

It seems clear that the bias described in this paper is caused by large, highly homogeneous, simple satellite blocks that affect read initiation, read extension, and base calling. However, what is the ultimate cause of these biases? The base-calling issue is the simplest to explain, as similar effects have been observed and explained before (Tan et al. 2022). Nanopore sequencing relies on detecting alterations in electrical conductance as single-stranded DNA traverses through a protein nanopore. Since the pore accommodates ∼6 nucleotides at a time, the raw signal reflects the joint effects of several bases and must be deconvoluted during base-calling. If a repeat monomer has a length close to 6 bp, it may confound the base-calling algorithm by producing minimal changes in electrical conductance. (Tan et al. 2022) found that this happened with the telomeric repeats (TTAGGG)n of several organisms ( *e.g.*, humans) and demonstrated that fine-tuning the Nanopore base-calling models improved accuracy in telomeric regions. A similar approach could improve the sequencing of *Drosophila* satellites, but this would not solve the major problem, which is the near absence of reads initiating within satellite blocks (Table 2). This is the most relevant question, and we address it in the next section.

### Why do reads seldom start within satellites?

The striking similarity between the PacBio and Nanopore coverage profiles (Fig. 1) suggests that the same underlying mechanism is responsible for low coverage in both technologies. Despite their fundamental differences - PacBio relies on a DNA polymerase and measures the fluorescence of modified DNA precursors while they are being incorporated into a nascent chain, whereas Nanopore measures the electrical condutance as native DNA passes through a membrane pore - one shared component stands out: T4 DNA ligase, which is used to glue sequencing adaptors in both platforms.

Evaluation of the possible role of T4 ligase is more complex than it might look at first sight. The enzyme exhibits different behaviors depending on the type of ligation (blunt ligation vs. cohesive-end ligation vs. nick ligation), and PacBio uses blunt-end ligation for gluing the SMRTbell adaptor to the target DNA (https://www.pacb.com/wp-content/uploads/2015/09/Guide-Pacific-Biosciences-Template-Preparation-and-Sequencing.pdf), whereas Nanopore uses a 1bp T/A overhang ligation (SQK-LSK114; https://nanoporetech.com/document/genomic-dna-by-ligation-sqk-lsk114). Some general patterns emerge from T4 ligase studies. Bauer et al. (2017) found T4 ligase to be among the least biased ligases, though human ligase III performed even better in this regard. They also reported that the T4 ligase has a preference for AT/TA over GC/CG for blunt end ligation, whereas Bilotti et al. (2022) found that GC is favored in cohesive end ligations. Additionally, ligation efficiency varies significantly even among substrates with the same GC content (Fig. 1 in Bilotti et al. 2022). Given these complexities, it seems difficult to derive from the T4 ligase properties an explanation for the satellite-induced bias we observed. But perhaps the main difficulty of such a hypothesis is that all known preferences of T4 ligase are short-range (*e.g.*, blunt end ligation of AT/TA vs GC/CG ends), whereas the effect of satellite DNA on TGS seems to obligatorily involve a much broader scale. For example, several satellites such as (AAAC)_n_ and (AAAAG)_n_ that we found near missing exons contain AT and GC base pairs, and hence should at least partially satisfy T4 ligase requirements, and yet they seldom were used as read initiation points (Table 2). The observation that only highly homogeneous satellite blocks exhibit these effects further suggests that the mechanism at play extends beyond a few base pairs.

Another way to investigate the role of the DNA ligase is to use protocols that replace it (at least in the main step of adaptor-target DNA ligation) with the transposase Tn5. This alternative method is available in the Nanopore Rapid Ligation Kit and, more recently, in the PacBio HiFi LILAP protocol (Jia et al. 2024). Fortunately, Jia et al. (2024) tested their LILAP method on *D. melanogaster* (iso-1 males), allowing us to examine its effect on the missing exons. As shown in Supplemental Fig. 8, the sequencing bias persists in LILAP reads, albeit to a lesser extent. This reduced bias is expected due to the smaller average size of LILAP HiFi reads (∼5kb; see comment about Illumina a few paragraphs above). One must also consider that Tn5 has its own sequence biases, which favor GC-rich regions (Kia et al. 2017), and that the LILAP protocol still includes a T4 ligase step to close the nick after the Tn5-mediated transposition. Hence, while DNA ligase remains a possible explanation for the missing exons, the limited available data does not support it as the primary cause.

Another possible explanation is that satellite DNA might be more resistant to shearing, reducing the number of reads that initiate in these regions. Indeed, Illumina sequencing bias seems to stem largely from DNA fragmentation biases, as sonication or nebulization (two commonly used fragmentation methods) preferentially induce breaks in the middle of CG dinucleotides (Poptsova et al. 2014, and references cited therein). Library preparation for TGS is always very gentle, which may facilitate a DNA fragmentation bias. It also occurred to us that most protocols for library preparation of TGS use a size selection buffer that selectively precipitates large DNA fragments; it is quite possible that satellite DNA does not behave as “normal” DNA in this respect and gets lost along with the “missing exons”.

While we focused more on the characterization and mechanistic aspects of the “missing exons”, there are other interesting (and more directly biological) questions. The strong biases caused by satellite DNA must be related to its poorly known properties *in vitro* and possibly *in vivo*. The data shown in Fig. 3 allow us to start looking at the sequence level at these mysterious regions of the *Drosophila* Y-chromosome in which protein-coding exons are embedded in huge blocks of intronic satellite DNA (Reugels et al. 2000), and yet are properly transcribed and spliced (Fingerhut et al. 2024). A complete assembly of the *Drosophila* Y is bound to shed light on some of these mysteries.

The missing exons phenomenon has obvious relevance for genomics and sequence technology; it most likely is not unique to *Drosophila*, and it is probably a matter of time before other cases of “missing exons” or unclosable gaps in assemblies are discovered or recognized. We hope this study stimulates both further investigations into its ultimate cause and improvements in sequence technology, and that eventually a T2T assembly of the *Drosophila* Y chromosome become feasible.

## Methods

### Raw reads

The raw reads used in this work are listed in Table S1. All datasets were obtained from adult *D. melanogaster* males of the reference strain *iso-1*. For the Illumina dataset, we extracted DNA from 40 freshly collected iso-1 males using the Promega “Wizard Genomic DNA Purification Kit” (cat # A1120), following the manufacturer’s recommendations. Library preparation and sequencing were performed at Macrogen (Korea), using the Illumina TruSeq Nano DNA PCR-free library protocol, 151 bp paired-end, with a 350 bp insert size.

### Genome assemblies

#### Illumina

The Illumina raw reads were first processed using TrimGalore (version 0.6.6) with Phred 33, a minimum read length of 77, and a stringency value of 4. To reduce the original ∼550× coverage to approximately 100×, reads were randomly subsampled using *seqtk* (Li 2012) before genome assembly. Assembly was performed using SPAdes version 3.15.3 (Bankevich et al. 2012) with default parameters. The final assembly was cleaned from contaminants using the *FCS-GX* program (Astashyn et al. 2024).

#### Nanopore

We started from the 400× dataset of Kim et al. (2024). We found that perhaps due to excess coverage a better assembly was obtained by removing reads shorter than 50 kb. This resulted in ∼84× coverage. The genome was then assembled using Flye (version 2.9.3) with the --nano-hq option. Following assembly, contaminants were removed using the *FCS-GX* program (Astashyn et al. 2024).

### Assembly of “missing exons” contigs

Normal-coverage regions of the Y chromosome were successfully assembled into large contigs using Nanopore reads (Kim et al. 2024) and Flye (Kolmogorov et al. 2019), as described above. However, all low-coverage regions were absent from this assembly, which was how we initially identified them. To reconstruct these missing exons, we used a targeted assembly approach. First, we used a blastN search using the CDS of each “missing exon” as the query and the Nanopore reads as the database and pulled all matching reads (we found 12 to 18 reads for each region). This procedure reduced assembly complexity by including only reads from the region of interest. We initially attempted to assemble each region separately, using Canu (Koren et al. 2017) and Flye (Kolmogorov et al. 2019), with very poor results (small contigs or no contig at all; it must be added that these tools were not designed for local assembly of highly repetitive regions, under shallow coverage). Using *minimap* / *miniasm* (Li 2016) and after trial and error on several parameters (including the undocumented parameter -S, which skips the last steps of *miniasm*), we eventually succeeded in assembling all “missing exon” regions into contigs ranging from 12 kb to 102 kb. Another helpful procedure was to remove before the assembly the reads that clearly are quimeric or rearranged (*e.g.*, Supplemental Fig. S10). Finally, as *miniasm* does not have a consensus step, we performed it using *racon* (Vaser et al. 2017). These steps are detailed in Supplemental Methods. The polished assembly of the “missing exons” (which should be considered draft assemblies) is available at https://github.com/bernardo1963/missing_exons. We also deposited there the sequences containing the normal coverage exons along with ∼ 50kb of flanking sequence on each side; these were used in comparisons with the low coverage regions (*e.g.*, Fig. 3). These sequences were extracted from the Flye whole genome assembly mentioned above.

### Statistical procedures

Most statistical tests were performed using SYSTAT 13 (nested ANOVA), or custom *Python* scripts based on the *statistics* and *scipy.stats* libraries (scripts are available at https://github.com/bernardo1963/missing_exons). For the Anderson-Darling test, we could not find a program or library that allows for a user-specified null hypothesis. To address this, we implemented the Anderson-Darling statistic in *Python* (allowing for a user-specified null hypothesis). We obtained the *P*-value by calling a modified version of the program *AnDarl.c* presented in (Marsaglia and Marsaglia 2004). These procedures are implemented in the *missingExon_stat_1jan2025.py* script, which is available at https://github.com/bernardo1963/missing_exons.

### Detection of repetitive sequences

We ran *censor* (Kohany et al. 2006) locally in order to detect and classify repetitive sequences present in the raw reads and assembled contigs. We found that several simple satellite repeats that are abundant in the “missing exons” contigs ( *e.g.*, (AATATAT)_n_ ) were not detected by *censor* due to their absence in its internal reference library (file *smprep.ref*). We fixed this problem by replacing the *smprep.ref* file with a complete, non-redundant list of all possible satellites up to 8 bp (file *satellite_8_pass2.fasta*, available at https://github.com/bernardo1963/missing_exons). In order to facilitate the joint annotation of repetitive sequences and the Y-linked exons (Fig. 3), we added to *censor* a custom library containing all Y-linked exons studied in this paper (file Ycdsrep.ref). We then ran censor with the following parameters:-lib Ycds -lib dro -lib inv -mode norm -bprm ’-filter=none’ -show_simple

### Read coverage estimation

For the Nanopore dataset, reads smaller than 1kb were excluded. This filtered dataset was then processed to remove adapter sequences using *porechop_abi* (Bonenfant et al. 2023) with the settings --ab_initio --no_split (we did this because adaptor sequences would interefere with read start point analysis; *e.g.*, Table 2). We obtained the read coverage data (*e.g.*, Fig. 1) by doing a blastN search of the CDS of the target genes against databases of sequencing reads (Nanopore, Illumina, *etc*.). The output was saved in tabular format (m8), and processed using a *python* custom script that reports the per base coverage and produces graphical representations of the data (*read_coverage_CDS_v4.py*, available at https://github.com/bernardo1963/missing_exons). We used WU-blast, but obtained essentially the same results when using NCBI-blast. We have not used *bwa* (Li and Durbin 2009) or similar read aligner programs because they all assume that the reference sequence contains all sequences present in the reads. This assumption was violated in our case, as we aligned genomic reads against a reference set of *Drosophila* CDS sequences rather than a complete genome assembly (which is not available for *Drosophila*). However, the results using read aligners were similar to ours (data not shown).

## Data access

The Illumina raw reads generated in this study have been submitted to the NCBI BioProject database (https://www.ncbi.nlm.nih.gov/bioproject/) under accession number PRJNA1227112. Scripts and data are included in the Supplemental Material and/or deposited in GitHub at https://github.com/bernardo1963/missing_exons.

## Acknowledgements

We thank Gustavo Khun, Cristiano Lazoski, Rodrigo Nunes, Thyago Vanderlinde, and our lab members for valuable suggestions during this work, and Jullien Flynn for help with the *D. virilis* data. This research was funded by FAPERJ - Fundação Carlos Chagas Filho de Amparo à Pesquisa do Estado do Rio de Janeiro, grant CNE2018, CNPq - Conselho Nacional de Desenvolvimento Científico e Tecnológico, grant INCT-EM, Wellcome Trust, grant 207486/Z/17/Z, to ABC. FU is supported by CAPES - Coordenação de Aperfeiçoamento de Pessoal de Nível Superior, Finance Code 001.

## Author Contributions

Conceptualization, methodology ABC; investigation, formal analysis, ABC and FU; data production: ABC and BYK; data curation: ABC, and FU; writing-original draft preparation, ABC; writing-review and editing, ABC, FU, BYK. funding: ABC. All authors have read and agreed to the submitted version of the manuscript.

